# The Geometric Sparse Matrix Completion Model for Predicting Drug Side effects

**DOI:** 10.1101/652412

**Authors:** Diego Galeano, Alberto Paccanaro

## Abstract

Pair-input associations for drug-side effects are obtained through expensive placebo-controlled experiments in human clinical trials. An important challenge in computational pharmacology is to predict missing associations given a few entries in the drug-side effect matrix, as these predictions can be used to direct further clinical trials. Here we introduce the Geometric Sparse Matrix Completion (GSMC) model for predicting drug side effects. Our high-rank matrix completion model learns non-negative sparse matrices of coefficients for drugs and side effects by imposing smoothness priors that exploit a set of pharmacological side information graphs, including information about drug chemical structures, drug interactions, molecular targets, and disease indications. Our learning algorithm is based on the diagonally rescaled gradient descend principle of non-negative matrix factorization. We prove that it converges to a globally optimal solution with a first-order rate of convergence. Experiments on large-scale side effect data from human clinical trials show that our method achieves better prediction performance than six state-of-the-art methods for side effect prediction while offering biological interpretability and favouring explainable predictions.

## 1 Introduction and Background

Drug side effects are a leading cause of morbidity and mortality in health care, with an annual cost of billions of dollars [1, 2, 3]. In this paper, we focus on the problem of predicting new unknown side effects for drugs for which a few experimentally determined side effects are already available. These computational predictions are important as they can be used after early-phase small-size human clinical trials, to set the direction of the risk assessment in later clinical trials, or after a drug has entered the market.

Several approaches have been proposed for predicting drug side effects (for reviews see [4, 5]) and can be roughly divided into two groups. The first group of methods exploits the network structure of the bipartite graph connecting drugs to side effects and networks built from drug or side effect side information. Cami et al. [6], for instance, built a bipartite network of drug side effects and extracted feature covariates from the network connectivity patterns to learn a Bernoulli expectation model based on multivariate logistic regression. Bean et al. [7] built a knowledge graph by connecting drugs, side effects, protein targets, and indications and then applied enrichment analysis to predict missing links in the network. Other network-based approaches include random walks and label propagation on side information networks [8, 9].

The second group of algorithms, proposed more recently, framed this problem as a matrix completion task using low-rank matrix decomposition techniques. Galeano and Paccanaro [10] used this type of model to predict missing associations in a binary matrix of drugs side effect associations. A similar approach was used by Zhang et al. [11], that also included smoothness constraints derived from drug side information. Li et al. [12] proposed an inductive matrix completion approach that integrates side information using kernel matrices of drugs and side effects.

In this paper, we cast the problem of drug side effect prediction as a *sparse high-rank* matrix completion problem. Our method is related to self-expressive models [13] that have recently been proposed as a framework for simultaneously clustering and completing high-dimensional data that lie in the union of low-dimensional subspaces. Self-expressive models can capture an underlying low-rank structure in a high-dimensional space or the union of low-rank structures leading to a full or high-rank structure [14, 15]. A self-expressive model represents each datapoint as a linear combination of a few other datapoints. Let *X* ∈ ℝ^*n*×*m*^ be the data matrix (each column is a datapoint) and let *C* ∈ ℝ^*m*×*m*^ be the coefficient matrix (each column is a coefficient vector). The goal of self-expressive model is to learn a matrix *C* such that *X* ≃ *XC* where *C* is sparse according to some sparsity function and diag(*C*) = 0 [14, 15]. Observe that the last constraint is needed to prevent the trivial solution of representing each datapoint with itself (*C* = *I*). Sparse linear method [16], proposed in the recommendation system literature, also shares the model assumption of self-expressive models.

### Contributions

We realized that the drug side effect matrix has a high-rank structure. We propose a novel high-rank sparse matrix completion model for predicting drug side effects. Extensive experiments on human clinical trials data show that our method outperforms existing state-of-the-art approaches in drug side effect prediction.

Our model is informative of the biology underlying drug activity: the learned (non-negative) sparse matrices of coefficients for drugs and side effects make explicit the similarities between drug activities at the molecular and phenotypic level. We show that these learned matrices of coefficients can be used for predicting the shared drug clinical activity, targets of drugs, and even the anatomical/physiological relationships between side effect phenotypes.

Our work is inspired by self-expressive models, but it differs from them as we assume that our data matrix is *fully* – rather than partially – observed while its entries are *noisy*. Our model incorporates structure into the learned matrices by exploiting side information graphs derived from the network structure of known relationships among row and column elements.

We prove that our multiplicative learning algorithm, which does not require to set a learning rate nor applying projection functions to guaranteed non-negative constraints, convergences to a globally optimal solution point with a first-order convergence rate. And unlike non-convex matrix decomposition models proposed previously for the side effect prediction problem [10, 11, 12], these theoretical guarantees of convergence imply the *reproducibility* of the solutions under arbitrary initializations: a desirable property for biological interpretation.

## 2 The Geometric Sparse Matrix Completion (GSMC) model

Let us denote our drug side effect matrix for *n* drugs and *m* side effects with the binary matrix *X* ∈ ℝ^*n*×*m*^ where *X*_*ij*_ = 1 if drug *i* is associated with side effect *j*, or 0 if the association is unreported. There are three main characteristics of *X*, which will need to be taken into consideration to build an effective algorithm. First, *X* is sparse (density ∼ 7%, see section 4); second, side effects have a long-tail distribution [17], which means that few side effects are responsible for the high proportion of entries in *X*; and third, unreported associations (zeros in *X*) have high uncertainty [17]. The last point stems from the fact that, typically, safety datasets report only observed pair-input associations. Consequently, a zero value represent the uncertain fact that either the drug does not cause the side effect, or that it does, but it could not be detected.

The analysis of our data matrix *X* reveals that the matrix has a high-rank (see section 4). Therefore, we cast the problem of side effect prediction as a sparse high-rank matrix completion problem for *X*. The goal of our Geometric Sparse Matrix Completion (GSMC) model is to *learn* two sparse matrices of coefficients, one for the row elements (*R* ∈ ℝ^*n*×*n*^) and one for the column elements (*C* ∈ ℝ^*m*×*m*^).The data matrix *X* is then approximated by:

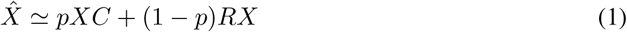

where *p* ∈ [0, 1] is a hyperparameter that controls the balance between the row (drug) and column (side effect) contributions. In the sequel, we shall refer to the first part of the GSMC model *XC*, as GSMC-c, and to the second part, *RX*, as GSMC-r. Two cost functions, *𝒬* _*c*_(*C*) and *𝒬* _*r*_(*R*), that takes into account side information for drugs and side effects are minimize with respect to *C* and *R*, respectively:

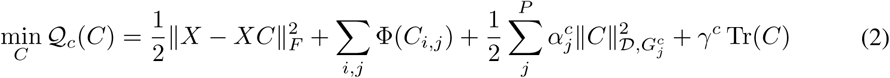

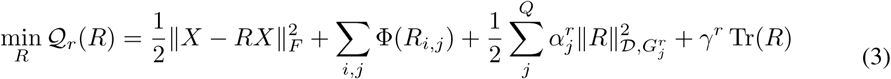

subject to the non-negative constraints *C, R* ≥ 0.

where ‖·‖_*F*_ is the Frobenius norm, Φ(·) is a sparsity function, and 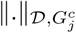 and 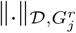 are the Dirichlet norms defined on *P* graphs 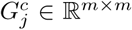, representing side information for side effects, and *Q* graphs 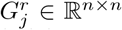, representing side information for drugs. *C* and *R* in Equation (1) are learned by minimizing Equations (2) and (3), respectively. In the following, we provide the rationale behind (2) only, as the same applies to (3).

The first term in Equation (2) is the *self-representation constraint*, which aims at learning a matrix of coefficients *C* such that *XC* is a good reconstruction of the original matrix *X* — as in self-expressive models, GSMC-c represents datapoints as a linear combination of other datapoints. The second term is the *sparsity constraint*, which uses the sparsity function 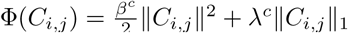^1^ – parameterized by constant values *β*^*c*^, *λ*^*c*^ *>* 0 – to favour sparse coefficients in the solution. The fourth term is the *null-diagonal constraint*, which has the important role of preventing the trivial solution *C* = *I* by imposing diag(*C*) = 0. This is achieved through a regularized trace operator *γ*^*c*^ Tr(*C*), whose parameter *γ*^*c*^ ≫0 does not need to be set by cross validation – the theoretical lower bounds for *γ*^*c*^ are provided in section 3.

Our model is called *geometric* due to the third term in Equation (3), the *smoothness constraint*, which incorporates structure into the sparse coefficient matrix *C*. This is achieved by adding smoothness priors from multiple weighted graphs that encode side information about the columns. Let us call one of these graph *G*^*c*^ ∈ ℝ^*m*×*m*^ (each node represents a side effect). Ideally, nearby points in *G*^*c*^ should have similar coefficients in *C*, which can be obtained by minimizing:

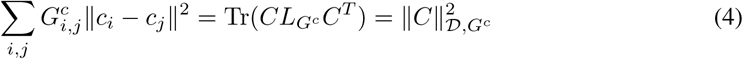

where *c*_*i*_ and *c*_*j*_ represent column vectors of *C*, 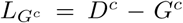 is the graph Laplacian, and 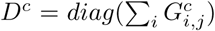 is a diagonal matrix. Extending this formulation to multiple graphs 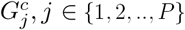 we obtain the third term in Equation (2): ^2^

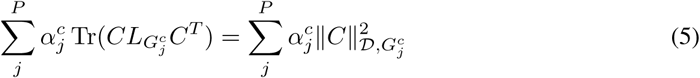

where the constant values 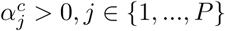 weigh the relative importance of each graph.

Finally, following [18], we impose *non-negative constraints* on *C*, as these constraints lead to more interpretable model since they allow only for additive combinations.

## 3 The Multiplicative Learning Algorithm

To minimize Equations (2) and (3) subject to the non-negative constraints *R,C* ≥ 0, we developed efficient multiplicative algorithms inspired by the diagonally rescaled principle of non-negative matrix factorization [18, 19]. The algorithm consists in iteratively applying the following multiplicative update rules:

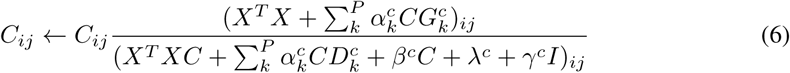

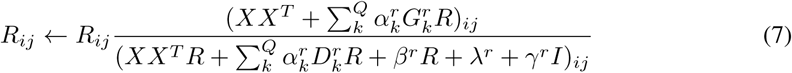

In the following, we shall prove that the algorithm in Eq. (6) converges to a solution; that the cost function *𝒬* _*c*_(*C*) is convex, and therefore the solution found is the global optimum; and that the speed of convergence is first-order. Finally we provide a lower bound for the value *γ*^*c*^. Proofs for Eq. (7) are similar and omitted here for brevity.

### Lemma 1.

*The cost function 𝒬*_*c*_(*C*) *in Equation (2) is convex in C.*

*Proof Sketch.* We need to prove that the Hessian is a positive semi-definite (PSD) matrix. That is, for a non-zero vector *h* ∈ ℝ^*m*^ the following condition is met *h*^*T*^ ∇^2^ *𝒬* _*c*_(*C*)*h* ≥ 0. The graph Laplacians are PSD by definition. The remaining terms in the Hessian (*X*^*T*^ *X* + *β*^*c*^) are also PSD. Therefore, *𝒬*_*c*_(*C*) is convex in *C*. See supplementary section S5 for complete proof.

### Theorem 1

(Convergence). *The cost function 𝒬*_*c*_(*C*) *in Equation (2) converges to a global minimum under the multiplicative update rule in (6).*

*Proof.* We need to show that our algorithm satisfies the Karush-Khun-Tucker (KKT) complementary conditions, which are both necessary and sufficient conditions for a global solution point given the convexity of the cost function (lemma 1) [20, 21]. KKT require *C*_*i,j*_ ≥ 0 and (∇ *𝒬*_*c*_(*C*))_*ij*_*C*_*ij*_ = 0. The first condition holds with non-negative initialization of *C*. For the second condition, the gradient is: 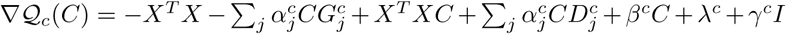, and according to the second KKT condition, at convergence *C* = *C*\* we have 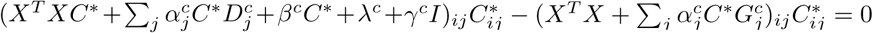, which is identical to (6). That is, the multiplicative rule converges to a global optima.

### Theorem 2

(Rate of convergence). *The multiplicative update rule in (6) has a first-order convergence.*

*Proof Sketch.* Following [20, 22], we can represent the updating algorithm as mapping *C*^*t*+1^ = ℳ(*C*^*t*^) with fixed point *C*\* = ℳ(*C*\*). Then, when *C*^*t*+1^ is near *C*\*, we have *C* ≃ ℳ(*C*\*) + ∇ ℳ(*C*)(*C* − *C*\*) subject to *C* ≥ 0, and thus ‖*C*^*t*+1^ − *C*\*‖ ≤ ‖∇ ℳ(*C*) ‖ · ‖*C*^*t*^ − *C*\*‖, with ‖∇ℳ(*C*) ‖ ≠ 0 almost surely. That is, the multiplicative update rule is a first-order algorithm.

### Theorem 3

(Lower bounds for the null-diagonal parameter *γ*^*c*^). *Let ϵ >* 0 *be the maximum tolerable value in diag(C)*, 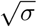 *the maximum initial value in diag(C), N*^*c*^ *the total number of iterations and L* = max_*i*_ *diag(X*^*T*^ *X). Then, γ*^*c*^ = *f* (*ϵ, N* ^*c*^) *is bounded by* 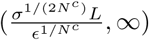.

*Proof.* Assuming that *γ*^*c*^ ≫ max_*i*_ diag 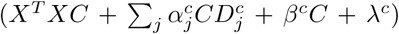 and that 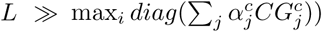, then at the *j*th iteration,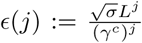. At convergence, *j* = *N*^*c*^, and 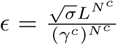, from which we can obtain the lower-bound for *γ*^*c*^. That is, to guarantee at most *ϵ* in diag(*C*), we need to set a 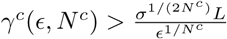. The upper bound is obtained when *ϵ* → 0, which causes *γ*^*c*^(*ϵ, p*) → *∞*. In practical applications, the upper bound is limited by machine precision.

The most expensive operation in (6) comes from the denominator term *X*^*T*^ *XC* for which *𝒪* (*N*^*c*^ ×*m*^3^) (where *N*^*c*^ is the total number of iterations). The overall complexity can be reduced by pre-computing the constant covariance matrix *X*^*T*^ *X* and the linear combination of graphs. A similar reasoning applies to (7), giving *𝒪* (*N*^*r*^ × *n*^3^). Algorithm 1 presents a Matlab pseudocode for solving GSMC-c that follows the NMF implementation guidelines in [23]: (i) *C*^*t*=0^ is sample from a uniform distribution in the interval 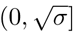; (ii) a small value *ε* ≃ 1 × 10^−16^ is added to the denominator to prevent division by zero. The stopping criteria for the algorithm is (i) when the number of iterations reaches `maxiter` or (ii) when the element-wise change 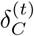 between *C*^(*t*+1)^ and *C*^(*t*)^ is smaller than a predefined tolerance `tolX`, with:

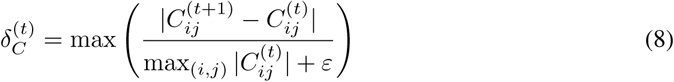

### ALGORITHM 1: GSMC-c

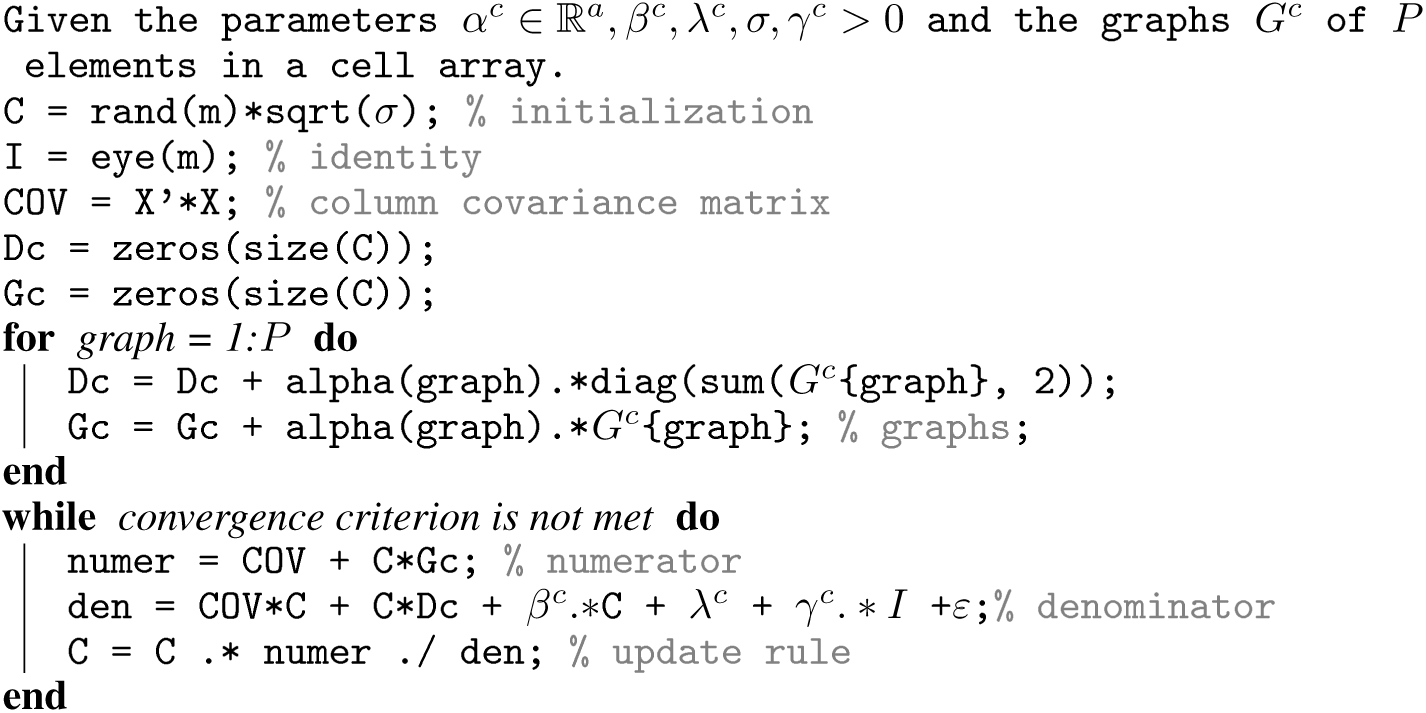

The algorithm to solve GSMC-r is similar and omitted for brevity. However, note that algorithm (1) can also be used to solve GSMC-r. This can be understood by considering that the GSMC-r model can be expressed as follow *RX* = (*X*^*T*^ *R*^*T*^)^*T*^ = (*Y A*)^*T*^ where *Y* = *X*^*T*^ and *A* = *R*^*T*^ and thus algorithm (1) can be used to solve *A* in *Ŷ* ≃ *Y A*.

## 4 Experimental Results

### Datasets

Drug side effects were extracted from the SIDER database [24, 25]. Our matrix *X* contains 75,542 known associations for 702 marketed drugs (rows) and 1,525 distinct side effect terms (columns) (7.06% density). Each drug and each side effect has at least six known associations. A value *X*_*ij*_ = 1 if a drug *i* is known to be associated with side effect *j* or *X*_*ij*_ = 0 otherwise (see Table S1 for details about the datasets).

In order to build graphs representing side information for drugs, we assembled binary matrices describing drug target interactions (702 drugs×401 targets), drug indication associations (702 drugs×5,178 indications), drug-drug interactions (702 drugs×702 drugs) and SMILES fingerprints – these datasets were extracted from DrugBank [26] and the Comparative Toxicogenomics database [27]. We then built the graphs using the cosine similarity between the rows of: the drug target matrix (we shall call this graph DT); the drug indication matrix (DInd); the drug-drug interaction matrix (DDI). The chemical graph (Chem) was built using the 2D Tanimoto chemical similarity from the drugs SMILES fingerprints (see section S4 for details). For each graph, we set the main diagonal of the weighted adjacency matrix to zero. The distribution of similarity scores of each graph is shown in Fig. S1. In the experiments, we did not include any graphs representing side information for side effects.

### Experimental setting

Following previous approaches [6, 8, 10, 11, 12], we frame the side effect prediction problem as a binary classification problem. We applied ten-fold cross-validation, while optimizing the hyperparameters using an inner loop of five-fold cross-validation within each of the ten folds (nested cross-validation for model selection [28]). The performance of the classifier is measured using the area under the receiver operating curve (AUROC) and the area under the precision-recall curve (AUPRC). We report the mean values of the ten folds for each metric (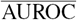 and 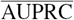). We compared the performance of our method against Matrix factorization (MF) [10], Inductive Matrix Completion (IMC)[12], Predictive PharmacoSafety Networks (PPNs) [6], Label propagation (LP)[8], Feature-derived graph regularized matrix factorization (FGRMF)[11], and side effect popularity (TopPop)[29]. While every algorithm used the drug side effect matrix *X*, only IMC, PPNs, LP and FGRMF could also make use of the drug side information graphs (see section S3 for a details for each model). Optimal hyperparameters for each model were optimized to maximize the 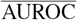 (see Table S5-S6). For GSMC, we optimize both models GSMC-c and GSMC-r independently. Then we set only the hyperparameter *p* using GSMC-c and GSMC-r with their obtained optimal hyperparameters. Datasets and code to reproduce the procedure are provided: (*This will be provided with the publication*).

### Performance evaluation

Table 1 summarizes the performance of the different methods. GSMC greatly outperforms the competitors both in terms of AUROC (by 1.3-19.7%) and in terms of AUPRC (by 4-30.9%). It is interesting to note that side effect popularity (TopPop) is highly predictive of drug side effects – this possibly reflects the fact that clinical reports are biased towards popular side effects such as headache or diarrhea [25]. The optimal value of *p* in GSMC was 0.45, indicating that although GSMC-c performs better than GSMC-r individually, the final model weighs the combination in favour of the latter, which includes side information about drugs. Our method also informs about the relative contribution of each side information: we found that molecular networks were weighted higher 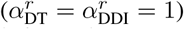, than networks containing chemical 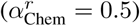 or phenotypic 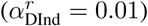 information. Importantly, we observed that the performance of our model is robust with respect to the setting of the model parameters *β*s and *λ*s (see the heatmaps in Fig. S4-S5).

**Table 1:** Performance comparison for drug side effect prediction

| Method | $\overline{\text{AUROC}} \pm \text{s.t.d.}$ | $\overline{\text{AUPRC}} \pm \text{s.t.d.}$ | Time(s)<sup>3</sup> |
| --- | --- | --- | --- |
| IMC [12] | $0.747 \pm 0.0113$ | $0.016 \pm 0.0011$ | $348.95 \pm 23.71$ |
| TopPop [29] | $0.827 \pm 0.0031$ | $0.071 \pm 0.0028$ | $0.010 \pm 0.0014$ |
| LP [8] | $0.888 \pm 0.0021$ | $0.126 \pm 0.0033$ | $0.018 \pm 0.0032$ |
| IMCZeros | $0.892 \pm 0.0045$ | $0.194 \pm 0.0100$ | $317.149 \pm 16.09$ |
| FGRMF [11] | $0.911 \pm 0.0029$ | $0.237 \pm 0.0059$ | $209.27 \pm 9.43$ |
| PPNs [6] | $0.923 \pm 0.0020$ | $0.208 \pm 0.0056$ | $186 \pm 5.91$ |
| MF [10] | $0.929 \pm 0.0019$ | $0.274 \pm 0.0071$ | $31.12 \pm 4.73$ |
| FGRMF-DDI [11] | $0.931 \pm 0.0020$ | $0.285 \pm 0.0075$ | $30.41 \pm 1.45$ |
| GSMC-r | $0.936 \pm 0.0014$ | $0.295 \pm 0.0073$ | $3.19 \pm 0.30$ |
| GSMC-c | $0.938 \pm 0.0023$ | $0.323 \pm 0.0024$ | $15.29 \pm 1.70$ |
| **GSMC** | **$0.944 \pm 0.0017$** | **$0.325 \pm 0.0063$** | **$17.82 \pm 1.95$** |

When comparing our method with competitor approaches, we found that a partial FGRMF [11] model based on the DDI graph only (FGRMF-DDI) performs better than the integrated model FGRMF – the fact that partial models could outperfom the integrated model had already been noted in the original publication. In the original publication [12], the IMC model was optimized using the observed entries only. Although matrix completion algorithms are predominantly based on this assumption [13, 15, 30, 31, 32, 33, 34, 35, 36], we found that taking into account the zeros can greatly improve the performance (we refer to this variant as IMCZeros in Table 1).

### High-rank structure of the drug side effects matrix

We verified that our 702 × 1, 525 drug side effect matrix *X* has a high rank – its value is 701^4^ (see the spectra in Fig. S2). We observed that the reconstructed matrices also preserve the high-rank structure, but with smooth filtering of the spectra, indicating that smaller singular values are important to model weaker regularities in the data (see Fig. S3).

### Biological interpretability

The effectiveness of our model at predicting the presence/absence of drug side effects prompted us to analyze whether the learned sparse matrices of coefficients are informative of the biology underlying drug activity. For these experiments, we trained the model using all the available data, fixed hyperparameters (*β*^*r*^ = 4, *λ*^*r*^ = 1, *β*^*c*^ = 2, *λ*^*c*^ = 0.5, *γ*^*c*^ = *γ*^*r*^ = 10^4^) and without side information graphs to avoid biases.

We first obtained a symmetrized version of the learned matrices *R* and *C*, defined as *𝒮* _*R*_ : = *R* + *R*^*T*^ and *𝒮* _*C*_ : = *C* + *C*^*T*^, respectively. Drug and side effect similarities were then defined as the cosine similarity between rows of *𝒮*_*R*_ and *𝒮*_*C*_, respectively. Drug clinical activity was defined using the Anatomical, Therapeutic and Chemical (ATC) taxonomy, a hierarchical organization of terms describing clinical activity where lower levels of the hierarchy contain more specific descriptors. Following the procedure in [17, 37, 38], we tested whether the similarity between two drugs was higher when they shared clinical activity.The evaluation was framed as a binary classification problem where the aim was to predict whether two drugs share an ATC category at different level of the taxonomy.

Figure 1a shows that our similarity is predictive of shared drug clinical activity. The predictions improve as we consider terms located lower in the ATC hierarchy (finer granularity) – this correctly reflects the fact that drug clinical responses become more similar as we move to lower (or more specific) levels of the ATC hierarchy. The figure also shows a comparison of the performance obtained for this problem with other methods used elsewhere [37, 39, 38]: Tanimoto chemical similarity and Jaccard side effect similarity (see section S4 for details). The fact that our similarity performs better than the Tanimoto chemical similarity in the chemical ATC subclass is quite remarkable, as in our model drugs are characterized only by noisy information about a few side effects, rather than exact knowledge of chemical structures.

**Figure 1:**
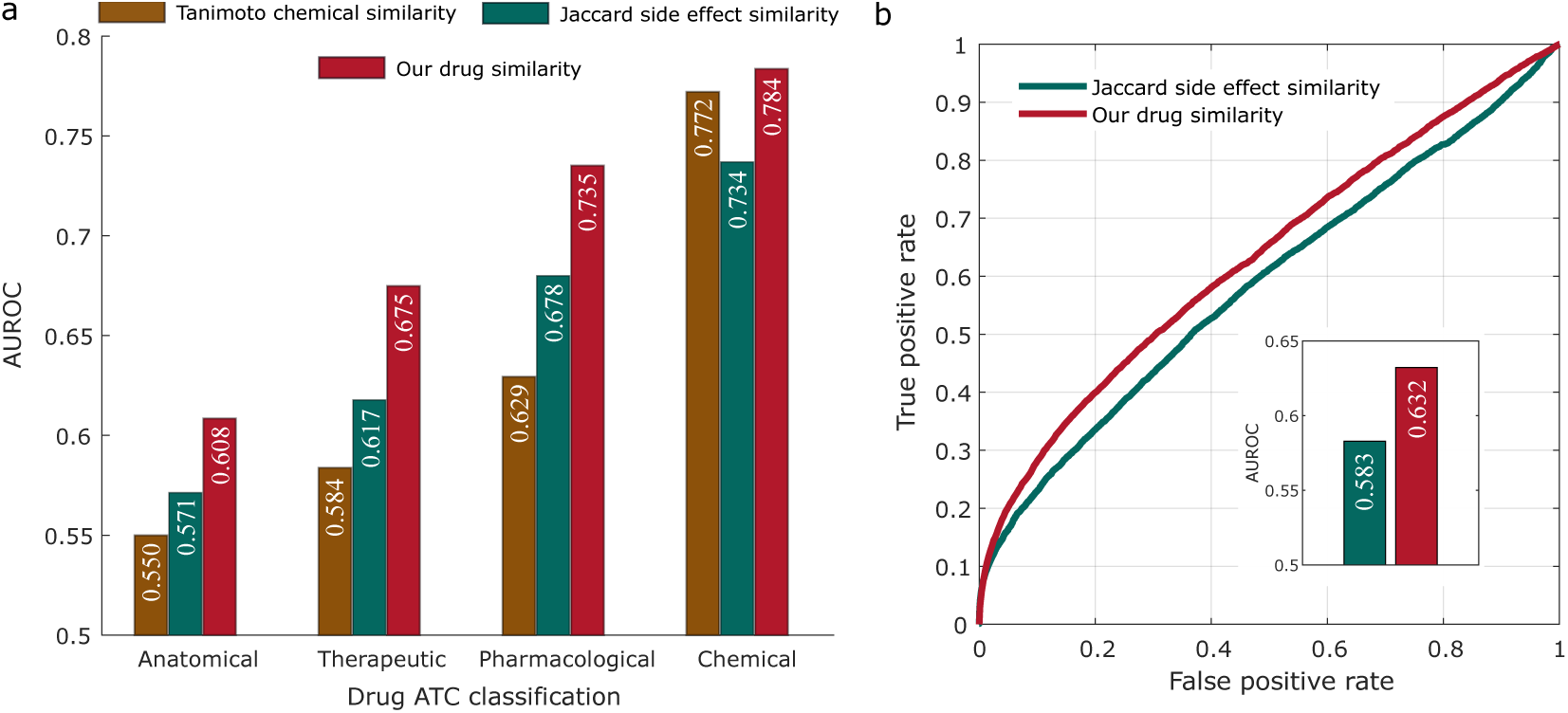
Our drug similarity captures drug clinical and molecular activity. **(a)** AUROC representing the performance of our drug similarity, side effect similarity (Jaccard) and Tanimoto chemical similarity at predicting whether a pair of drugs share Anatomical, Therapeutic and Chemical (ATC) category at each level of the ATC taxonomy. **(b)** ROC curve representing the performance of our drug similarity at predicting whether pairs of drugs share a target. *Inset* AUROC barplot.

Encouraged by these results, we decided to test whether our drug similarity could even be used for the prediction of shared drug targets. Having framed this as a binary classification problem, we found that our drug similarities are predictive of shared protein targets between drugs (see Figure 1b). Note that, drug side effect similarity had previously been found to be predictive of drug protein targets at molecular level [40, 38], but the fact that our similarity, that is built using the same data, works better, means that our model is able to exploit the information more effectively (4% AUROC improvement). Finally, we found that using the cosine similarity between the rows of *𝒮*_*R*_, instead of *𝒮*_*R*_directly, slightly improves the prediction performance – this is probably due to the fact that the cosine similarity is less noisy as it takes into account the similarity between all the neighbourhs of each drug. Fig. S6 presents the embedding of drugs in 3D based on *𝒮*_*R*_ that is obtained applying t-SNE [41] together with the heatmap of the mean inter- and intra-class similariy *𝒮*_*R*_ for each ATC anatomical classes.

Finally, we also analyzed the link between side effect similarities and the anatomy/physiology of the side effect phenotypes. Side effects were grouped based on their anatomical class according to MedDRA [42]. We found that similarities for two side effects tend to be higher when they are phenotypically related. Figure S7 shows that, in most cases, the side effect similarity within system organ classes (top level of the MedDRA hierarchy) is higher than the similarity between classes. Moreover, side effect similarity is predictive of shared MedDRA category at each of the different levels and predictions improve as we move to more specific terms in the MedDRA hierarchy.

## 5 Conclusion and Discussion

In this paper, we show that the drug side effect matrix has a high rank structure, and we presented a novel high-rank sparse matrix completion approach based on geometric multi-graph learning to predict side effects of drugs that outperforms state of the art models. To our knowledge, our work is the first that relies on the high-rank assumption to predict drug side effects. We envision the application of our geometric sparse matrix completion model to other problems in computational biology and pharmacology with similar high-rank structure.

An advantage of our method is that the predictions are *explainable* thanks to the non-negative constraints on the learned matrices. Fig. 2 shows an example using the GSMC-c model and Lindane, a drug that has been withdrawn from the market due to side effects that had gone unreported during clinical trials. Lindane is amongst the drugs with the smallest number of side effects in our dataset (1.5^th^ percentile) – only 10 side effects are present. Figure 2a shows the histogram of the values found in the row corresponding to Lindane in *XC*. Our model predicts that Lindane is likely to cause hypotension (the score is in the 98.8^th^ percentile) and indeed this side effect has been repeatedly reported [43, 44]. Figure 2b provides the rationale behind this prediction. The score for Lindane-hypothension is the sum of the (non-negative) coefficients in the column of *C* corresponding to hypotension for the 10 known side effects of Lindane. Notice how `seizures`, a condition normally associated to hypothension, explains 37.92% of the score strength. As illustrated in this example, an analysis of the non-negative coefficients learned by our model can potentially provide biological clues to generate medical and pharmacological hypothesis when assessing the safety of a drug.

**Figure 2:**
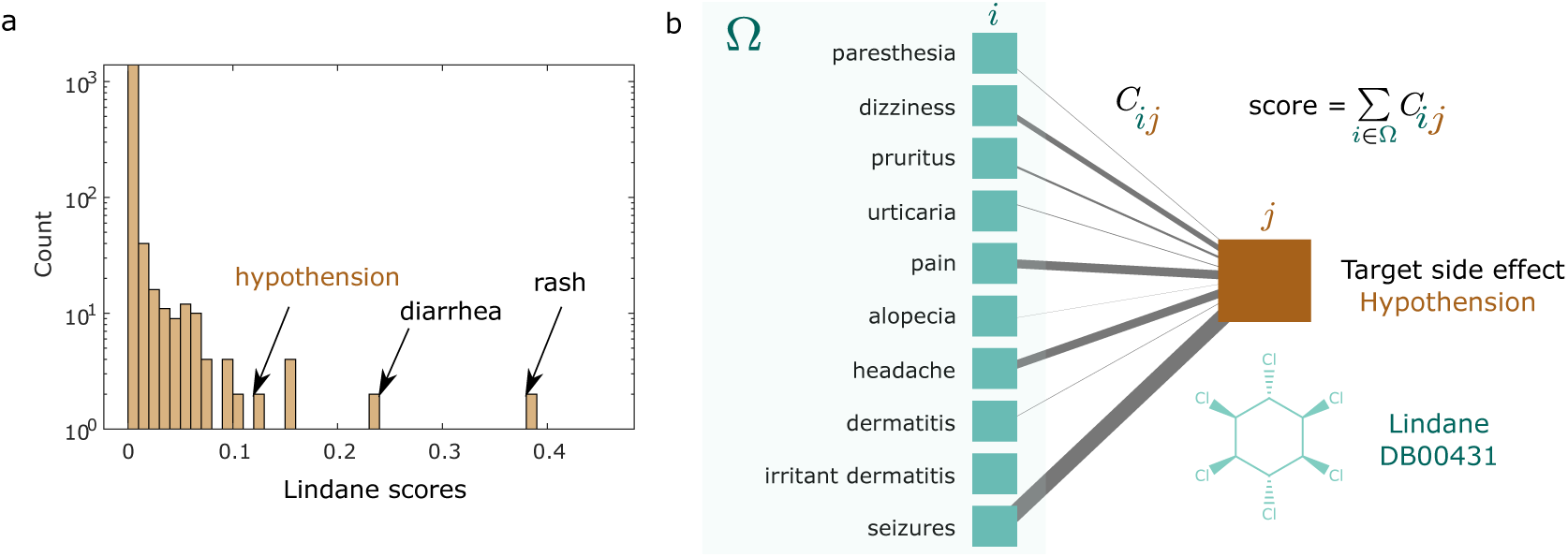
Example of explainable predictions for the withdrawn drug Lindane. **(a)**. Histogram of predicted scores for Lindane using GSMC-c; **(b)** Network diagram despicting how the model generates the predictions for a given target side effect under study. In the figure, Ω represents the set of known side effects indexed by *i*, and *j* is the target side effect. The thickness of the connections are proportional to the learned coefficients.

## Footnotes

1 This function is also known as the elastic-net regularization.

2 Note that for Equation (3), the graphs 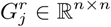 have a different number of nodes (each node represents a drug) and the Dirichlet norm is applied to the rows of 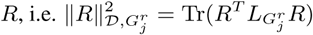.

3 Average time of running the algorithm in the ten fold cross-validation.

4 This was computed using the Matlab built-in function `rank`.

