## Supplementary Materials for "The Geometric Sparse Matrix Completion Model for Predicting Drug Side effects"

**S4 Biological interpretability of our model** **7**

**S5 Convexity of the cost function** **8**

**S6 Supplementary Figures** **9**

**S7 Supplementary Tables** **16**

**References** **18**

Preprint. Under review.

#### S1 Dataset details

Table S1: Public datasets

| dataset matrix | source | #rows | #cols | #nnzs | density |
|---|---|---|---|---|---|
| drug side effects | SIDER2 [12] | 702 drugs | 1,525 side effects | 75,542 | 7.06% |
| chemical similarity | DrugBank 3.0 [11] | 702 drugs | 702 drugs | 490,112 | 100% |
| protein targets | DrugBank 3.0 [11] | 702 drugs | 401 targets | 1,918 | 0.65% |
| drug-drug interactions | DrugBank 3.0 [11] | 702 drugs | 702 drugs | 10,024 | 2.03% |
| drug indications | CTD [7] | 702 drugs | 5,178 indications | 198,323 | 5.45% |

The columns #rows, #cols and #nnzs indicate the number of drugs, other association and non-zero entries, respectively. The density is the percentage of #nnzs from the total number of entries, i.e. density = #nnzs / (#drugs×#cols).

To study the biological interpretation of the model, we also obtained the drugs Anatomical, Therapeutic and Chemical (ATC) classification code from the World Health Organization (WHO) release 2018. For side effects, we obtained the Medical Dictionary for Regulatory Activities (MedDRA) classification of disorders (version 21.0).

#### S2 Computing infrastructure used and implementations

All the models were run in a small cluster with 32 physical cores and 100GB of RAM. Each processor is a Intel(R) Xeon(R) CPU E5-2683 v4 @ 2.10GHz. We implemented most of the algorithms in Matlab R2018a 64-bit. Training of model hyperparameters was executed in parallel using the Parallel Computing Toolbox version 6.12.

#### S3 Methods implementation and optimization

For completeness, in this section, we introduce each baseline method, the details about the implementation, optimization, and hyperparameters tunning. Partial results of the models (with each side graph) is also shown in some cases.

##### S3.1 Side effect popularity (TopPop)

Cremonesi et al. [6] used movie popularity as a baseline to assess the performance of movie recommendation system algorithms. We adopted the same baseline in our problem of drug side effect prediction. We were motivated by recent work that indicates that the distribution of drug side effects is similar to the distribution of movie datasets [10]. Given the data matrix $X \in \mathbb{R}^{n \times m}$ for $n$ drugs and $m$ side effects, TopPop assigns scores to drug side effect pairs $(i, j)$ as follow:

$$\text{TopPop}_{ij} = \sum_{i}^{n} X_{ij} \quad (1)$$

##### S3.2 Low-rank matrix factorization (MF)

The MF model [9] is based on the assumption that the binary matrix $X \in \mathbb{R}^{n \times m}$ for $n$ drugs and $m$ side effects of rank $k \ll min\{m, n\}$, can be expressed as the product of two matrices of rank-$k$, as follow:

$$X \approx PQ \quad (2)$$

where $P \in \mathbb{R}^{n \times k}$ and $Q \in \mathbb{R}^{k \times m}$ . In this model, $P$ represents the drug latent space and $Q$ the side effects latent space. The product $PQ$ is related to the probability that a given drug causes certain side effects. The matrices $P$ and $Q$ are the solution of the following optimization problem:

$$\min_{P,Q} \mathcal{L}(P, Q) = \min_{P,Q} \frac{1}{2}\|X - PQ\|_F^2 + \frac{\lambda}{2}(\|P\|_F^2 + \|Q\|_F^2) \quad (3)$$

where the term $\lambda(\|P\|_F^2 + \|Q\|_F^2)$ is an L2 regularization term with penalty $\lambda$, which is added to the objective function in order to prevent over-fitting, and $\|.\|_F$ is the Frobenius norm defined as the square root of the sum of the square of its elements. The problem is non-convex in both $P$ and $Q$. Gradient descent-based methods are required to approximate the solution. Thus, the derivatives of (3) are:

$$\frac{\partial \mathcal{L}(P,Q)}{\partial P} = -(X - PQ)Q^T + \lambda P \tag{4}$$

$$\frac{\partial \mathcal{L}(P,Q)}{\partial Q} = -P^T(X - PQ) + \lambda Q \tag{5}$$

**Optimization** Following the procedure presented in [9], we used conjugate gradient descent (CGD) and approximated line searches based on polynomial interpolation with Wolfe-Powel conditions to find the local minima in (3). We used the same Matlab implementation of the Rassmusen minimizer used by the authors[1].

**Hyperparameters tunning** Following the procedure presented in [9], we initialize $P$ and $Q$ as normally distributed random variables with small variance $\sigma^2 = 0.01$. We tuned both model parameters: the number of latent factors $k$ and the regularization penalty $\lambda$ in the grid: $k \in \{10, 30, 50, 70, 90, 100\}$ and $\lambda \in \{0.1, 1, 5, 10, 15, 20\}$.

##### S3.3 Predictive Pharmacosafety Networks (PPNs)

PPNs begins by creating a bipartite graph in which one set of nodes represent the drugs and the other set of nodes represent the side effects. The set of edges corresponds to the known drug-side effect associations. PPNs then computes covariate measures in this network to capture the structure of connections between drugs and side effects. The covariate $Z_s(u, j)$ quantify a relationship between a drug node $u$ and a side effect node $j$. The predictive model is based on the binary response variable $X_{uj}, u \in \{1, ..., n\}, j \in \{1, ..., m\}$, for $n$ drugs and $m$ side effects. $Y_{uj}$ denotes the presence or absence of drug-side effect associations. Using a multivariate logistic-regression model, the response is modeled as a Bernoulli random variable with the following expectation

$$\mathbb{E}[X_{uj}] = \frac{1}{1 + \exp\left(-\sum_s \beta_s Z_s(u, j)\right)} \tag{6}$$

where $\beta_s$ denotes the model parameter and $Z_s$ the model covariate. The optimization is performed by assuming independence between the responses $X_{ij}$ and performing model fitting by maximum likelihood. Several covariates are considered in the PPN model. The *degree covariates* are defined as follow;

$$\begin{aligned} Z_1(u, j) &= \text{deg}(u) \times \text{deg}(j) \\ Z_2(u, j) &= |\text{deg}(u) - \text{deg}(j)| \\ Z_3(u, j) &= \text{deg}(u) + \text{deg}(j) \\ Z_4(u, j) &= \frac{\text{deg}(u)}{\text{deg}(j)} \end{aligned} \tag{7}$$

Here, deg$(u)$ denotes the degree of node $u$. The degree product covariate, $Z_1(u, j)$, aimed to capture preferential attachment among high-degree drugs and side effects. The degree difference covariate, $Z_2(u, j)$, aimed to capture assortativity, i.e. whether high-degree drugs connects to high-degree side effects or to small-degree side effects. The *distance covariates* are also defined. These depends not

[1] `http://learning.eng.cam.ac.uk/carl/code/minimize/`

only on the nodes $(u, j)$ but on the sets of neighbor $N(u)$ and $N(j)$. Let $J(j, k)$ denotes the Jaccard similarity between the neighbors sets $N(j)$ and $N(k)$,

$$J(j,k) = \frac{|N(j) \cap N(k)|}{|N(j) \cup N(k)|} \tag{8}$$

The following Jaccard-based covariates quantify structural similarity between drug pairs and side effect pairs,

$$\begin{aligned} Z_5(u,j) &= \max_{k \in N(u)-\{j\}} \{J(j,k)\} \\ Z_6(u,j) &= \max_{k \in N(j)-\{u\}} \{J(u,k)\} \end{aligned} \tag{9}$$

Finally, Jaccard-based predictors based on Kullback-Leibler divergence ($\mathcal{K_L}$) between the overall distribution of similarities between a drug ($\bar{D}_{se}$) and the drugs in its local neighborhood ($D_{se}(u, j)$) or between side effects ($\bar{D}_{se}$) and side effects in its neighborhood ($D_{se}(u, j)$) are defined;

$$\begin{aligned} Z_7(u,j) &= \mathcal{K_L}(D_{se}(u,j), \bar{D}_{se}) \\ Z_8(u,j) &= \mathcal{K_L}(D_{drug}(u,j), \bar{D}_{drug}) \end{aligned} \tag{10}$$

We also include all the four drug graphs used in the paper, but following the procedure presented in [4] to include drug side information. The following drug side information covariates were also included:

$$\begin{aligned} Z_9(u,j) &= \min_{k \in N(j)-\{i\}} \{d_{chem}(u,k)\} \\ Z_{10}(u,j) &= \min_{k \in N(j)-\{i\}} \{d_{DDI}(u,k)\} \\ Z_{11}(u,j) &= \min_{k \in N(j)-\{i\}} \{d_{DT}(u,k)\} \\ Z_{12}(u,j) &= \min_{k \in N(j)-\{i\}} \{d_{DInd}(u,k)\} \end{aligned} \tag{11}$$

Each pairwise distance between drugs was built from the affinity graphs (distance = $1 - G$).

**Optimization** We use the Matlab built-in function `mnrfit` (link function `logit`) to learn the coefficients estimates ($\beta_s$) of the multinomial logistic regression of the nominal responses in $X$ on the predictors in $Z$.

**Hyperparameters tunning** PPNs does not require hyperparameters tunning.

##### S3.4 Inductive Matrix Completion (IMC)

IMC [14] is based on a matrix decomposition model with low-rank assumption that allows for the integration of additional information about drugs and side effects. Given a sample of observed entries $\Omega$ from a binary matrix of drug side effects $X \in \mathbb{R}^{n \times m}$ for $n$ drugs and $m$ side effects, the goal is to estimate the missing entries. IMC assumes that the matrix is low-rank $X \approx WH$, where $W \in \mathbb{R}^{n \times k}$ and $H \in \mathbb{R}^{k \times m}$. Let now consider a kernel similarity matrix for drugs $K_d \in \mathbb{R}^{n \times n}$ and a kernel similarity matrix for side effects $K_a \in \mathbb{R}^{m \times m}$. The goal is to learn $X$ using the observed entries $\Omega$, by minimizing the following loss:

$$\min_{W,H} \mathcal{L}(W,H) = \frac{1}{2}\|\Omega \circ (X - K_d W H K_a)\|_F^2 + \frac{\lambda}{2}(\|W\|_F^2 + \|H\|_F^2) \tag{12}$$

where $\Omega$ represents a projection for the observed entries in $X$, $\circ$ is the element-wise product between matrices, the term $\lambda(\|W\|_F^2 + \|H\|_F^2)$ is an L2 regularization term with penalty $\lambda$, which is added to the objective function in order to prevent over-fitting, and $\|.\|_F$ is the Frobenius norm defined as the square root of the sum of the square of its elements. The problem is non-convex in both $W$ and $H$. Gradient descent-based methods are required to approximate the solution. Thus, the derivatives of (12) are:

$$\frac{\partial \mathcal{L}(W,H)}{\partial W} = -K_d(X - \Omega \circ (K_d W H K_a))K_a H^T + \lambda W \tag{13}$$

$$\frac{\partial \mathcal{L}(W,H)}{\partial H} = -W^T K_d(X - \Omega \circ (K_d W H K_a))K_a + \lambda H \tag{14}$$

Kernels matrices were built following the procedure used in [14] using our dataset. We linearly combined the similarity matrices as follow:

$$K_d = \frac{1}{4}G_{chem} + \frac{1}{4}G_{DDI} + \frac{1}{4}G_{DT} + \frac{1}{4}G_{DInd} \tag{15}$$

where $G_{chem}, G_{DDI}, G_{DT}, G_{DInd} \in \mathbb{R}^{n\times n}$ are chemical similarity, drug interaction cosine similarity, drug target cosine similarity and drug indication cosine similarity, respectively. For side effects, the kernel $K_a$ was simply obtained by computing the cosine similarity between the columns in $X$ (for each training set).

**Optimization** We used conjugate gradient descent (CGD) and approximated line searches based on polynomial interpolation with Wolfe-Powel conditions to find the local minima in (12).

**Hyperparameters tunning** We initialize $P$ and $Q$ as normally distributed random variables with small variance $\sigma^2 = 0.01$. We tuned both model parameters: the number of latent factors $k$ and the regularization penalty $\lambda$ in the grid: $k \in \{10, 30, 40, 50, 70, 90, 100\}$ and $\lambda \in \{0.1, 1, 5, 10, 15, 20\}$.

##### S3.5 Feature-derived graph regularized matrix factorization (FGRMF)

The FGRMF [17] model relies on low-rank assumption but integrates drug side information using smoothness constrains on the drug latent representations. Given a binary association matrix $X \in \mathbb{R}^{n\times m}$ for $n$ drugs and $m$ side effects, the goal of FGRMR is to approximate $X$ by the product of two low-rank matrices $\hat{X} \approx WH$, $W \in \mathbb{R}^{n\times k}$ and $H \in \mathbb{R}^{k\times m}$, for an integer $k \ll \min(n, m)$. FGRMR uses a two step learning to integrate multiple side information. In the first step, it optimizes a low-rank model using one side information graph. In the second step, uses a logistic regression model to integrate the optimal models obtained independently. In detail:

1 **Low-rank model with side information**. Given a side information graph for drugs $G \in \mathbb{R}^{n\times n}$, the goal of FGRMF is to minimize the following objective:

$$\min_{W,H} \mathcal{L}(W,H) = \min_{W,H} \frac{1}{2}\|X - WH\|_F^2 + \frac{\lambda}{2}(\|W\|_F^2 + \|H\|_F^2) + \frac{\alpha}{2}\|W\|_{\mathcal{D},G}^2 \tag{16}$$

where $L_G = D - G$ is the graph Laplacian, $D = diag(\sum_i G_{i,j})$ and $\|.\|_{\mathcal{D},\mathcal{G}}$ is the Dirichlet semi-norm of the graph $G$.

2 **Integration using LR**. Given independent scores for each side information $\hat{X}_{\text{chem}}, \hat{X}_{\text{DDI}}, \hat{X}_{\text{DT}}$ and $\hat{X}_{\text{DInd}}$, FGRMF generates a probability scores ($h_\theta(\hat{X})$) by optimizing the following model:

$$h_\theta = \frac{1}{1 + \exp\left(-(\theta_0 + \theta_1\hat{X}_{\text{chem}} + \theta_2\hat{X}_{\text{DDI}} + \theta_3\hat{X}_{\text{DT}} + \theta_4\hat{X}_{\text{DInd}})\right)} \tag{17}$$

where the weights $\theta_j$ is the weight learned for each independent model.

**Optimization** We used conjugate gradient descent (CGD) and approximated line searches based on polynomial interpolation with Wolfe-Powel conditions to find the local minima in equation (16). To optimize equation (17), we use the Matlab built-in function `mnrfit` (link function `logit`) to learn the coefficients estimates ($\theta_j$).

**Hyperparameters tunning** We initialize $W$ and $H$ as normally distributed random variables with small variance $\sigma^2 = 0.01$. We tuned the three model parameters: the number of latent factors $k$, the regularization penalty $\lambda$ and the penalty $\alpha$ in the grid: $k \in \{20, 30, 40, 50, 60, 70, 80, 90, 100\}$ and $\lambda \in \{1, 5, 10, 15, 20\}$ and $\alpha \in \{0.1, 1, 2, 3, 4, 5\}$. The results for individual FGRMF models in shown in Table S2. Interestingly, partial models performed better than the LR integration, which was consistent with the results found by the authors in [17].

Table S2: Performance of partial FGRMF models

| Model | $\overline{\text{AUROC}}\pm$ s.t.d. | $\overline{\text{AUPRC}}\pm$ s.t.d. | params $(\lambda, k, \alpha)$ |
|---|---|---|---|
| FGRMF-Chem | $0.929 \pm 0.0021$ | $0.278 \pm 0.0076$ | 1,70,1 |
| FGRMF-DDI | $0.931 \pm 0.0020$ | $0.285 \pm 0.0075$ | 10,70,0.1 |
| FGRMF-DT | $0.930 \pm 0.0021$ | $0.277 \pm 0.0075$ | 10,70,1 |
| FGRMF-DInd | $0.928 \pm 0.0020$ | $0.270 \pm 0.0053$ | 1,70,1 |
| Logistic regression (LR) | $0.911 \pm 0.0029$ | $0.237 \pm 0.0059$ | - |

The column `params` indicate the optimal hyperparameters obtained in cross validation.

##### S3.6 Label propagation using consistency method (LP)

One of the first large-scale methods to predict drug side effects was proposed in [1]. Atias et al. tested their method for de-novo prediction, that is, to predict side effects for drugs assuming no known initial associations for the drugs (by removal of entire rows in the matrix). We tested label propagation using the Zhou et al.[18] consistency method for our problem using our datasets.

We tested four different LP models, one based on side effect similarity graphs and the others based on drug side information graphs. We describe briefly the consistency method for the second case. Given a drug-drug similarity network $G \in \mathbb{R}^{n\times n}$, and initial labels in a matrix $Y \in \mathbb{R}^{n\times m}$ for $n$ drugs and $m$ side effects. $Y_{ij} = 1$ if drug $i$ is known associated to side effect $j$ or $Y_{ij} = 0$ otherwise.

The algorithm consists in the following steps [1, 18]:

1. Given the affinity graph $G$, set $G_{ii} = 0$.
2. Construct the matrix $S = G \oslash \sqrt{P^T P}$, where $\oslash$ indicates element-wise division and $P = \sum_i G_{ij}$ is the sum of the rows of $G$.
3. Compute the scores $F = (I - \alpha S)^{-1} Y$, where $I \in \mathbb{R}^{n\times n}$ is the identity matrix and $\alpha$ is a free-parameter that trades between the neighborhood network information and its initial label information.

Results of the partial models are shown in Table S3. We could not use the side graphs DT and DDI because they were too sparse to perform the label propagation in those individual graphs (singular matrix in the inversion process). For the model LP-all, we use a graph that was the linear combination of all the available graphs:

$$G_{all} = \frac{1}{4}G_{chem} + \frac{1}{4}G_{DDI} + \frac{1}{4}G_{DT} + \frac{1}{4}G_{DInd} \tag{18}$$

Table S3: Performance of individual LP models

| Model | Graph similarity measure | $\overline{\text{AUROC}}\pm$ s.t.d. | $\overline{\text{AUPRC}}\pm$ s.t.d. | params $(\alpha)$ |
|---|---|---|---|---|
| LP-SE | Side effects Jaccard | $0.888 \pm 0.0021$ | $0.126 \pm 0.0033$ | 0.1 |
| LP-chem | Tanimoto ($G_{chem}$) | $0.834 \pm 0.0033$ | $0.075 \pm 0.0030$ | 0.001 |
| LP-DInd | Cosine ($G_{DInd}$) | $0.840 \pm 0.0030$ | $0.078 \pm 0.0028$ | 0.001 |
| LP-all | $G_{all}$ | $0.845 \pm 0.0031$ | $0.086 \pm 0.0035$ | 0.001 |

The column `params` indicate the optimal hyperparameters obtained in cross validation.

**Optimization** We used the close form of $F$ to obtained the scores for drug side effects.

**Hyperparameters Tunning** We tuned $\alpha$ from 0.001 to 0.9 in steps of 0.01 (90 values).

##### S3.7 Geometric Sparse Matrix Completion (GSMC)

The Matlab code used to run our experiments for GSMC-c (and GSMC-r) is provided in section **??**.

**Optimization** We followed the recommended guidelines used to implement non-negative matrix factorization (NMF) in [2]. $R$ and $C$ were initialized with weights from a uniform distribution between $(0, \sqrt{\sigma}]$ for a $\sigma = 0.01$. The maximum number of iterations was set to 100 and `tolX` = 0.01. With this `tolX`, convergence occurs in roughly 50 iterations. To set $\gamma^c$, $\gamma^r$, using our theoretical bounds, we observed that, for GSMC-c $L^c = \max_i \text{diag}(X^T X) = 602$, and for GSMC-r, $L^r = \max_i \text{diag}(XX^T) = 644$. Thus, a $\gamma^c = \gamma^r = 10^4$ is enough to obtain an $\epsilon \approx 9.54 \times 10^{-63}$. Therefore, $\gamma^c$ and $\gamma^r$ were set to $10^4$ for all the experiments.

**Hyperparameters tunning** For the partial GSMC-c model we tuned the parameters in the following grid $\beta^c \in \{0, 0.1, 0.5, 1, 2, 3, 4, 5, 10, 20\}$, $\lambda^c \in \{0, 0.5, 1, 2, 3, 4, 5\}$. For the partial GSMC-r model that integrates side information, we reduced the grid due to the larger number of possible combinations: $\beta^r \in \{1, 2, 3, 4, 5, 10\}$, $\lambda^r \in \{0.1, 0.5, 1\}$ and $\alpha^r \in \{0.01, 0.1, 0.5, 1\}$. Finally, to train the GSMC model (the linear combination of GSMC-c and GSMC-r), we tuned only $p \in \{0, 0.01, 0.02, ..., 1\}$ while setting the following hyperparameters for the partial models: GSMC-c ($\beta^c = 1, \lambda^c = 0.5$) and GSMC-r ($\beta^r = 2, \lambda^r = 0.5, \alpha^r_{chem} = 0.5, \alpha^r_{DDI} = 1, \alpha^r_{DT} = 1, \alpha^r_{DInd} = 0.01$). These settings were based on the optimal values during model selection of each partial model.

#### S4 Biological interpretability of our model

In the main manuscript, we show that our learned sparse matrices of coefficients have a biological meaning. Here we present in detail the similarities used: Jaccard side effect similarity, 2D Tanimoto chemical similarity and our similarities based on $C$ and $R$.

Baseline similarities were defined following the procedure in [5, 10, 16]:

- **Jaccard side effect similarity**. Given two vectors $x_a, x_b \in \mathbb{R}^{m \times 1}$ describing the presence/absence of $m$ side effects for drug $a$ and $b$, respectively, the Jaccard similarity between the vectors is defined as follow:

$$\mathcal{J}(x_1, x_2) = \frac{\#(x_1 \cap x_2)}{\#(x_1 \cup x_2)} \tag{19}$$

  where $\#$ indicates number in the intersection and union of sets. The Jaccard similarity is bounded $0 \leq \mathcal{J}(x_1, x_2) \leq 1$.

- **2D Tanimoto chemical similarity**. Given two vectors describing the hash binary fingerprints $h_a, h_b \in \mathbb{R}^{\#\text{bits} \times 1}$ extracted from drugs chemical structures (SMILES representation), the Tanimoto similarity is defined as follow:

$$\mathcal{T}(h_a, h_b) = \frac{\#(h_a \cap h_b)}{\#(h_a \cup h_b)} \tag{20}$$

  here $\#$ indicates number in the intersection and union of sets. The Tanimoto similarity is the Jaccard similarity of the binary fingerprints. Tanimoto similarity was computed directly from the SMILES representation of the drugs using RDKit in python [13].

- **Our drug similarity**. Given that our model's sparse matrix of coefficients is not symmetric ($R \neq R^T$), we first obtained the similarity between drugs as follow [8]:

$$\mathcal{S}_R = R + R^T \tag{21}$$

  where $\mathcal{S}_R \in \mathbb{R}^{n \times n}$ is a symmetric similarity matrix for the $n$ drugs [2]. To account for the neighborhood information of a drug, the final similarity used in the main manuscript is the cosine distance of the rows in $\mathcal{S}_R$.

[2]Note that our similarity matrix does not contain self-similarity (because the optimization enforces diag($R$) = 0 to prevent the trivial solution), which can be simply added by adding a identity matrix to $\mathcal{S}_R$.

- **Our side effect similarity**. Similarly, to obtain side effect similarities we computed the cosine similarity of the rows of $\mathcal{S}_C = C + C^T$.

Table S4: AUROC performance at predicting drug clinical activity

| ATC category | Chemical | Side effect | $\mathcal{S}_R$ | CosSIM of $\mathcal{S}_R$ | AUROC improvement (%) |
|---|---|---|---|---|---|
| Anatomical (Level 1) | 0.550 | 0.571 | 0.599 | **0.608** | 3.70 |
| Therapeutic (Level 2) | 0.584 | 0.617 | 0.66 | **0.675** | 5.80 |
| Pharmacological (Level 3) | 0.629 | 0.678 | 0.723 | **0.735** | 5.70 |
| Chemical (Level 4) | 0.772 | 0.734 | 0.771 | **0.784** | 3.80 |

Best values are shown in bold. AUROC improvement percentages are computed based on the best AUROC using chemical or side effect similarities.

#### S5 Convexity of the cost function

Given that GSMC cost functions are convex, then the Karush-Khun-Tucker (KKT) equations are both necessary and sufficient conditions for a global solution point [3]. Here we formalize this by proving that the GSMC-c objective function is *convex*, therefore, there are not spurious local minimum in the algorithms to solve GSMC-c and GSMC-r. To learn $C$, we minimize:

$$\min_C \mathcal{Q}_c(C) = \frac{1}{2}\|X - XC\|_F^2 + \sum_{i,j} \Phi(C_{i,j}) + \frac{1}{2}\sum_j^P \alpha_j^c \|C\|^2_{\mathcal{D},G_j^c} + \gamma^c \operatorname{Tr}(C) \tag{22}$$

We shall now prove the following theorem.

**Theorem 1** (Convexity)**.** *For a real matrix $X$, the objective function $\mathcal{Q}_c(C)$ is convex in $C$.*

*Proof.* We need to show that the Hessian of $\mathcal{Q}_c(C)$ is a positive semi-definite (PSD) matrix. The Hessian of $C$ is the second derivative of $\mathcal{Q}_c(C)$, that is:

$$\nabla^2 \mathcal{Q}_c(C) = X^T X + \beta^c + \sum_j \alpha_j^c L_{G_j^c}$$

where $L_{G_j^c} = D_j^c - G_j^c$ is the graph Laplacian. The Hessian is PSD iif for a non-zero column vector $h \in R^m$ of real numbers the scalar $h^T \nabla^2 \mathcal{Q}_c(C) h \geq 0$, i.e.,

$$\begin{aligned} h^T(X^T X + \beta^c + \sum_j \alpha_j^c L_{G_j^c})h &\geq 0 \\ h^T X^T X h + \beta h^T h + \sum_j h^T \alpha_j^c L_{G_j^c} h &\geq 0 \\ \| Xh \|^2 + \beta \| h \|^2 + \sum_j \alpha_j^c \| y_j \|^2 &\geq 0 \quad \forall h, y, \beta^c > 0 \end{aligned} \tag{23}$$

In the last step, we use the fact that the Laplacian $L_{G_j^c}$ is PSD and thus can be diagonalized as follow:

$$L_{G_j^c} = U_j^T \Lambda_j U_j = U_j^T \Lambda_j^{\frac{1}{2}} \Lambda_j^{\frac{1}{2}} U = (\Lambda_j^{\frac{1}{2}} U)^T (\Lambda_j^{\frac{1}{2}} U)$$

Letting $y_j = \Lambda_j^{\frac{1}{2}} U$, we see that:

$$h^T \alpha_j L_{G_j^c} h = \alpha_j y_j^T y_j = \alpha_j \| y_j \|^2 \geq 0$$

That is, the GSMC-c objective function is convex in $C$. □

### S6 Supplementary Figures

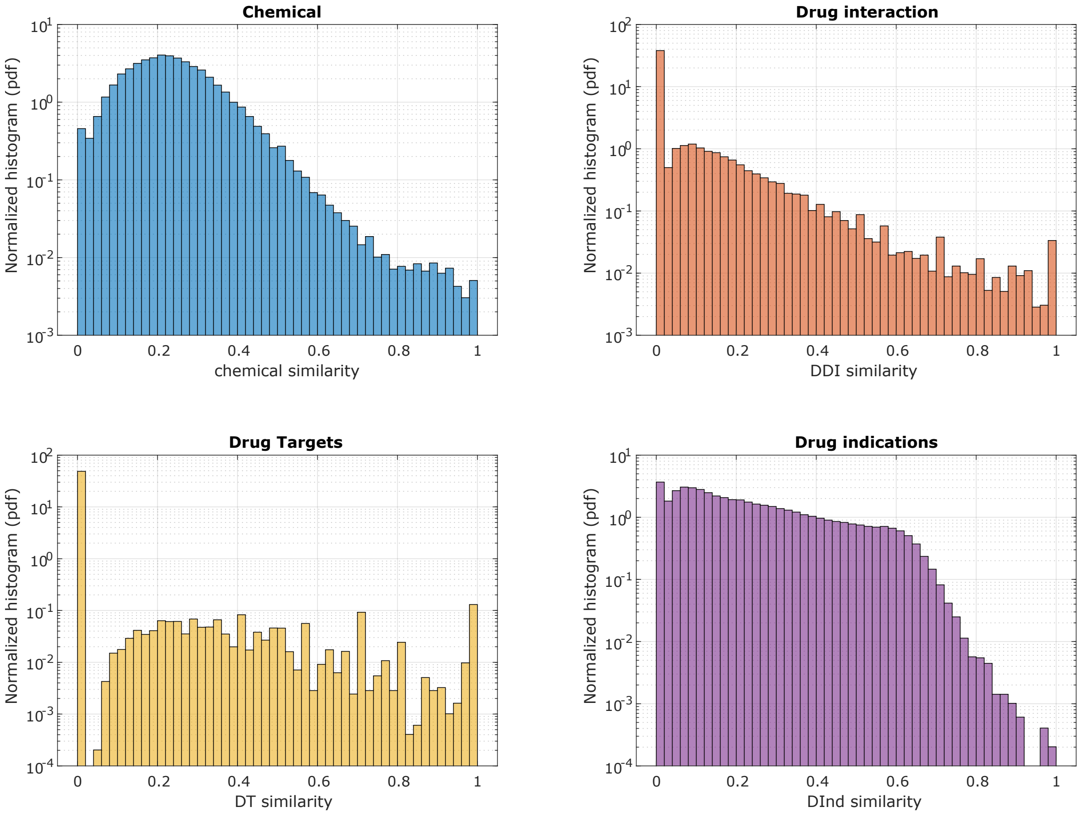

Figure S1: **Normalized histogram of similarities used as graph side information for drugs**: chemical similarity (blue), drug interaction (orange), drug targets (yellow) and drug indications (purple). All the similarities are bounded in the interval $[0, 1]$.

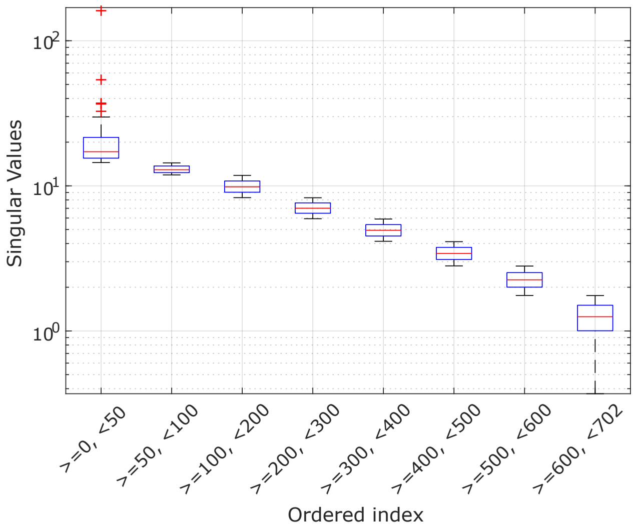

Figure S2: **High rank structure of drug side effects**. Boxplots of singular values of the data matrix $X$ of drug side effects. Singular values were group according to their ordered index. The drug side effects data matrix has a high-rank: `rank(X) = 701`. And even the distribution of singular values 600th to 702th (the group with smaller singular values) rank significantly higher than zero (Wilcoxon Signed Rank Significance, $p < 1.83 \times 10^{-18}$).

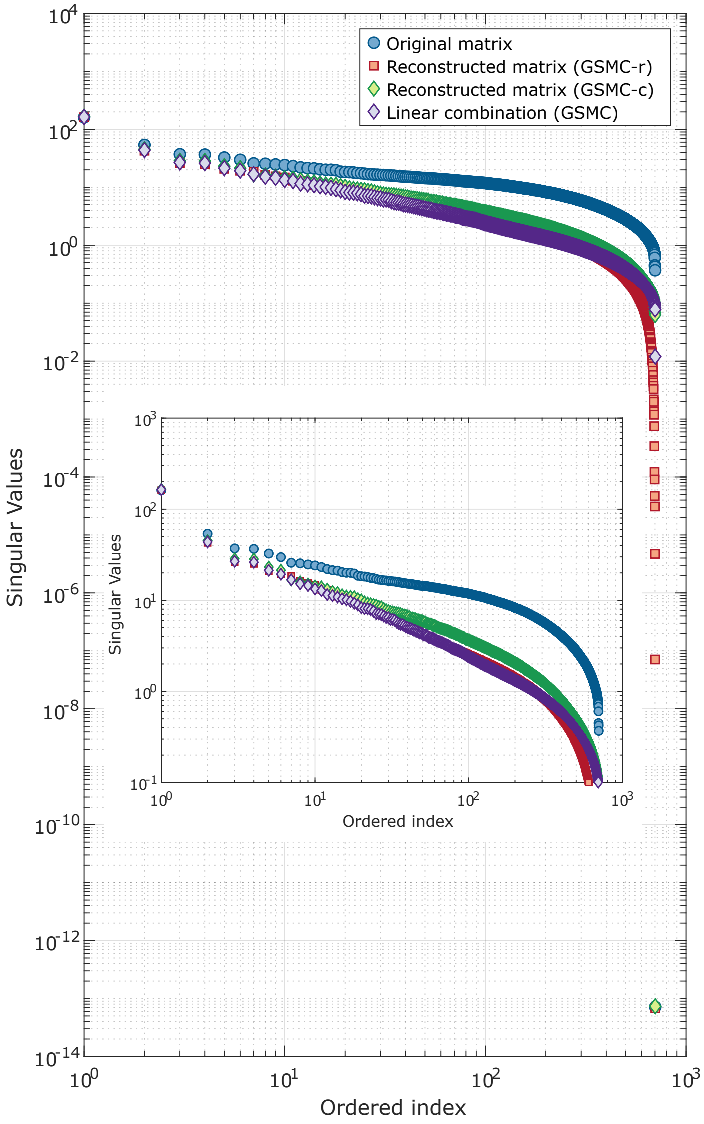

Figure S3: **Smooth filtering of the spectra of singular values**. Singular values of the original matrix $X$ and the reconstructed matrices $RX$, $XC$ and $\hat{X} \simeq \frac{1}{2}RX + \frac{1}{2}XC$. For this experiment, we did not consider side information graphs. The models GSMC-r and GSMC-c performs a smooth spectral filtering (de-noising). The density of the reconstructed matrix by GSMC-r is 47.74% ($R$ has a density of 19.09%) and 48.83% by GSMC-c ($C$ has a density of 5.58%). The threshold we used to calculate the densities was 0.01 for the reconstructed matrices and $1 \times 10^{-4}$ for the sparse matrices. *Inset*. Zoom into a region of the spectra.

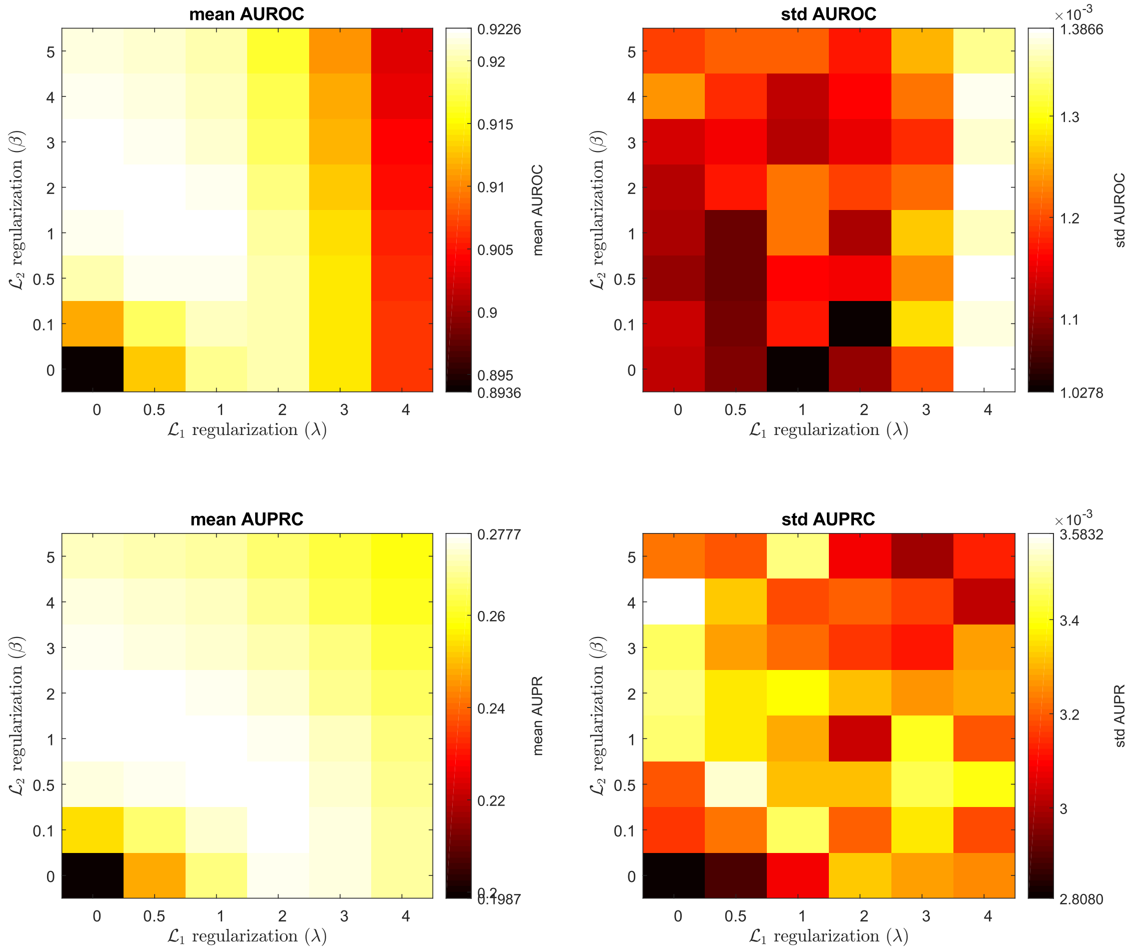

Figure S4: **Heatmaps of the average performance of GSMC-c during model selection across the five-fold cross-validation in the validation sets**. The performance is consistent across folds (small standard deviation) and it is not very sensitive to the setting of the model hyper parameters. Optimal performance can also be achieved by only using $\beta^c > 0$ (with $\lambda^c = 0$).

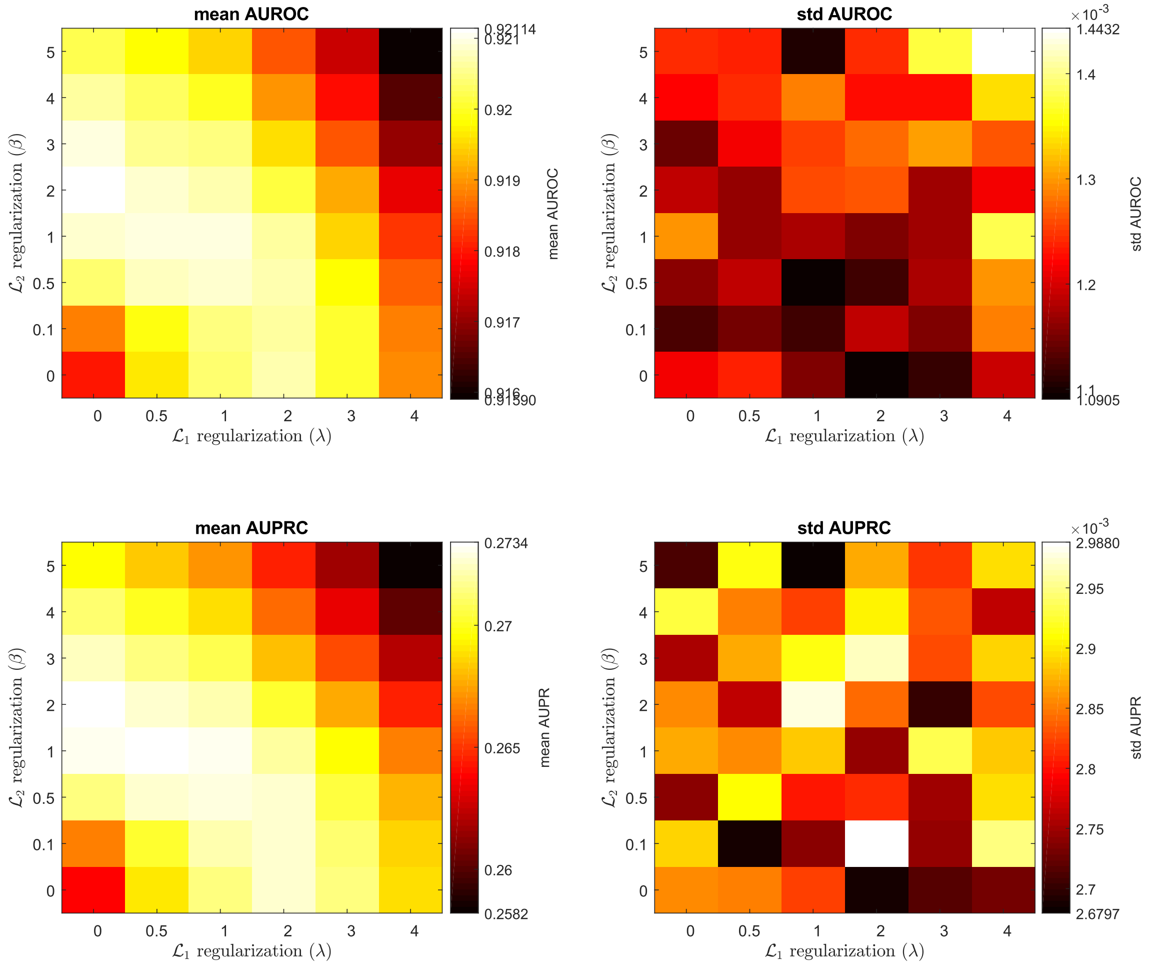

Figure S5: **Heatmaps of the average performance of GSMC-r (without side graph) during model selection across the five-fold cross-validation in the validation sets**. The performance is consistent across folds (small standard deviation) and it is not very sensitive to the setting of the model hyper parameters. Optimal performance can also be achieved by only using $\beta^r > 0$ (with $\lambda^r = 0$).

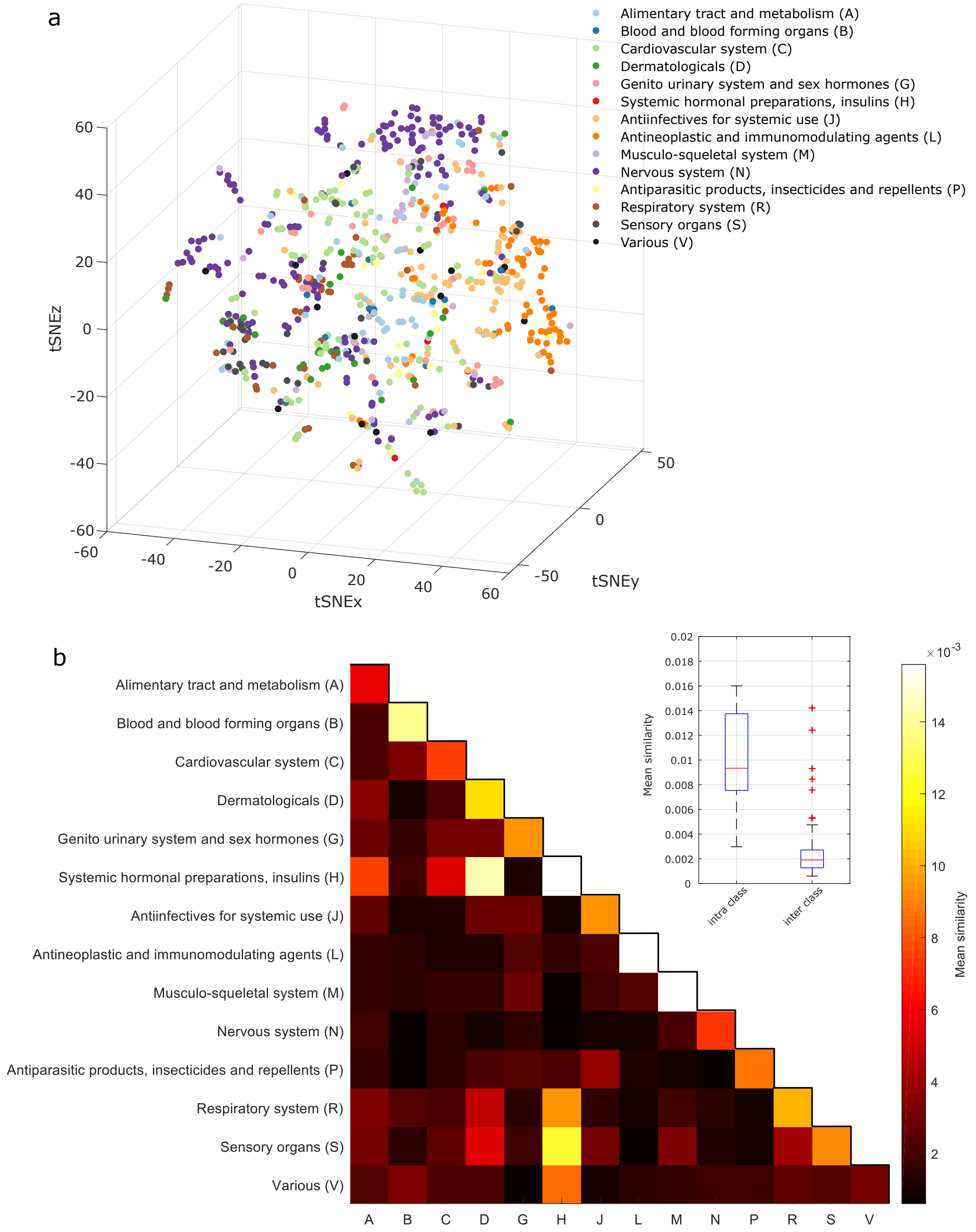

Figure S6: **Drug sparse matrix of coefficients similarity captures drug clinical activity (a)** Embedding of drugs in 3D space using t-SNE [15]. Each point represents a drug. Colours are assigned based on their anatomical category. Distance between points is related to the cosine distance of the drug clustering similarity $\mathcal{S}_R = R + R^T$. **(b)** Heatmap of mean drug similarities $\mathcal{S}_R$ per anatomical class. Each $(x, y)$ tile represents, for each main Anatomical, Therapeutic and Chemical (ATC) drug category, the mean similarity of drug pairs where one drug belong to category $x$ and the other to category $y$. The value ranges from $3 \times 10^-4$ (Musculo squeletal system - Systemic Hormonal and Preparations) to $0.0078$ (Musculo squeletal system-Musculo squeletal system). The colours range between the minimum mean similarity and $0.0156$, with all values above $0.0156$ (In the diagonal: $0.0921$ (H), $0.0160$ (M)) set to $0.0156$. *Inset*: the average intra-class similarity is significantly higher than the average interclass similarity (t-test Significance, $p < 7.12 \times 10^{-13}$).

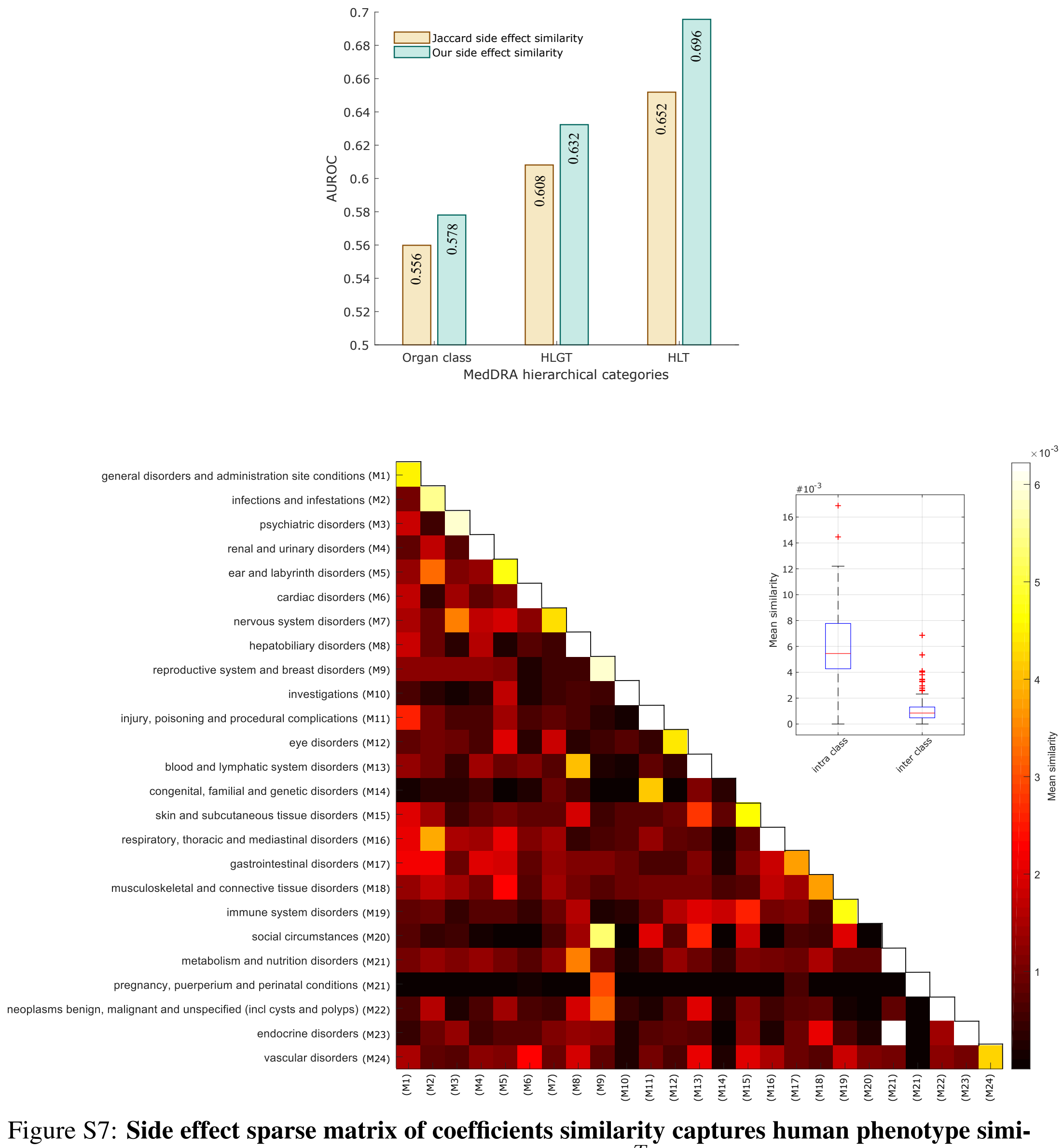

Figure S7: **Side effect sparse matrix of coefficients similarity captures human phenotype similarity (Top)** Ability of our side effect similarity ($\mathcal{S}_C = C + C^T$) and the Jaccard side effect similarity to predict whether two side effects belong to the MedDRA class at different levels of the hierarchy. **(Bottom)** Heatmap of mean side effect similarities $\mathcal{S}_C$ per organ class. Each ($x$, $y$) tile represents, for each main MedDRA organ class, the mean similarity of side effect pairs where one side effect belong to category $x$ and the other to category $y$. The value ranges from $1.29 \times 10^{-24}$ (M21 - M14) to 0.017 (M8-M8). The colours range between the minimum mean similarity and 0.0062, with all values above 0.0062 (In the diagonal: 0.0075 (M4), 0.0098 (M6), 0.0169 (M8), 0.010 (M10), 0.0067 (M11), 0.014 (M13),0.0062 (M16),0.0065( M21), 0.00863 (M23),0.012 (M24); off-diagonal:0.00686 (M24-M21)) set to 0.0062. *Inset*: the average intra-class similarity is significantly higher than the average interclass similarity (t-test Significance, $p < 7.14 \times 10^{-81}$).

#### S7 Supplementary Tables

Table S5: Optimal hyper-parameters for each fold in cross-validation for the different methods

| Fold number | IMC [14] $(k, \lambda)$ | IMCZeros $(k, \lambda)$ | MF [9] $(k, \lambda)$ | FGRMF-chem [17] $(k, \lambda, \alpha)$ | FGRMF-DDI [17] $(k, \lambda, \alpha)$ | FGRMF-DT [17] $(k, \lambda, \alpha)$ | FGRMF-DInd [17] $(k, \lambda, \alpha)$ |
|---|---|---|---|---|---|---|---|
| 1 | **50,10** | **20,20** | **70,15** | **70,1,1** | **70,10,0.1** | **70,10,1** | 20,5,0.1 |
| 2 | 30,10 | 20,20 | 80,15 | 70,1,1 | 70,10,0.1 | 70,10,1 | **70,1,1** |
| 3 | 40,10 | 20,20 | 90,15 | 70,1,1 | 70,10,0.1 | 70,10,1 | 20,5,0.1 |
| 4 | 50,10 | 20,20 | 90,15 | 70,1,1 | 70,10,0.1 | 70,10,1 | 70,1,1 |
| 5 | 40,10 | 20,20 | 100,15 | 70,1,1 | 70,10,0.1 | 70,10,1 | 70,1,1 |
| 6 | 20,10 | 20,20 | 100,15 | 70,1,1 | 70,10,0.1 | 70,10,1 | 20,5,0.1 |
| 7 | 10,20 | 20,20 | 80,15 | 70,1,1 | 70,10,0.1 | 70,10,1 | 70,1,1 |
| 8 | 20,10 | 20,20 | 70,15 | 70,1,1 | 70,10,0.1 | 70,10,1 | 70,1,1 |
| 9 | 50,10 | 15,20 | 70,15 | 70,1,1 | 70,10,0.1 | 70,10,1 | 70,1,1 |
| 10 | 30,5 | 15,20 | 50,15 | 70,1,1 | 70,10,0.1 | 70,10,1 | 70,1,1 |

Bold indicates the most consistent hyparameter values in the cross-validation (by majority rule).

Table S6: Optimal hyper-parameters for each fold in cross-validation for the different GSMC models

| Fold number | GSMC-c $(\beta, \lambda)$ | GSMC-r (no graph) $(\beta, \lambda)$ | GSMC-r (chem) $(\beta, \lambda, \alpha)$ | GSMC-r (DDI) $(\beta, \lambda, \alpha)$ | GSMC-r (DT) $(\beta, \lambda, \alpha)$ | GSMC-r (DInd) $(\beta, \lambda, \alpha)$ | GSMC-r (all) $(\beta, \lambda, \alpha_{chem}, \alpha_{DDI}, \alpha_{DT}, \alpha_{DInd}))$ | GSMC $p$ |
|---|---|---|---|---|---|---|---|---|
| 1 | 2,0 | 1,0 | 3,1,0.5 | 20,1,1 | 5,1,0.7 | 2,1,0.5 | 10,1,0.1,1,1,0.1 | 0.44 |
| 2 | 1,0.5 | 2,0 | 0.1,0.5,0.5 | 20,1,2 | 5,2,1 | 2,1,0.5 | 4,0.5,0.5,0.5,1,0.01 | 0.42 |
| 3 | 1,1 | 1,0.5 | 3,0.5,0.5 | 20,1,1 | 4,2,0.7 | 2,0.5,0.5 | 2,0.5,0.5,0.5,0.5,0.1 | 0.43 |
| 4 | 3,0 | 2,0.5 | 4,0.5,0.5 | 20,0.5,2 | 5,1,1 | 4,0.5,0.5 | 3,0.5,0.5,0.5,0.5,0.01 | 0.42 |
| 5 | 1,0.5 | 1,1 | 0,0.5,0.5 | 20,0.5,2 | 4,0.5,1 | 3,0.5,0.5 | 5,1,0.1,1,1,0.1 | 0.45 |
| 6 | 2,0 | 2,0.5 | 2,0.5,0.5 | 20,1,2 | 5,0.5,1 | 5,0.5,0.5 | 1,1,0.1,0.5,1,0.1 | 0.45 |
| 7 | 1,0.5 | 3,0 | 0,0.5,0.7 | 10,1,2 | 3,2,0.7 | 0.5,1,0.5 | 10,0.5,0.1,0.5,1,0.1 | 0.45 |
| 8 | 1,0.5 | 2,0 | 4,1,0.5 | 20,1,2 | 4,1,1 | 1,1,0.5 | 5,1,0.1,0.5,1,0.1 | 0.48 |
| 9 | 3,0 | 2,1 | 0.5,1,0.5 | 20,0.5,1 | 5,2,1 | 5,1,0.5 | 10,1,0.1,0.5,1,0.1 | 0.46 |
| 10 | 3,0 | 1,0.5 | 4,0.5,0.5 | 20,1,2 | 5,1,0.7 | 5,0.5,0.5 | 10,0.1,0.1,1,1,0.1 | 0.46 |

For all the cases, $\gamma = 10^4$. For the final GSMC model, only $p$ was trained, the following parameters were set based on the individual model's performances: GSMC-c $(\beta = 1, \lambda = 0.5)$ and GSMC-r (*all*) $(\beta = 2, \lambda = 0.5, \alpha_{chem} = 0.5, \alpha_{DDI} = 1, \alpha_{DT} = 1, \alpha_{DInd} = 0.01))$.

[3] Stephen Boyd and Lieven Vandenberghe. *Convex optimization*. Cambridge university press, 2004.

[10] Diego Galeano and Alberto Paccanaro. Predicting the frequency of drug side effects. *bioRxiv*, 2019.
